# A modelling framework to estimate vector dispersal and disease spread in an important agricultural pathosystem

**DOI:** 10.1101/2024.12.16.628743

**Authors:** Katharine F. Preedy, Louise Mc Namara, Md Munir Mostafiz, Maximilian Schughart, Lawrence E. Bramham, Daniel J. Leybourne

## Abstract

Cereals are some of the most agriculturally important crops grown worldwide. Cereal aphids are small sap-feeding insects that infest cereals and vector devastating cereal pathogens, principally the yellow dwarf viruses, with severe infection reducing yields by 20 *−* 80%. Currently, farmers are advised to apply insecticide if a single cereal aphid is found within the crop. However, this is an oversimplification of the biology and ecology of the vector-virus system: Each aphid species can transmit a range of yellow dwarf virus species with variable efficiency, and intra-species diversity within a vector species (genotype) can influence this further. Accounting for this variation in decision-making processes has the potential to help with developing bespoke management options based on the composition of the local vector-virus population, and represents a novel avenue that could be explored to develop more sustainable management strategies. Here we describe a stage-structured model that can estimate vector dispersal and disease spread. We use data for the two main cereal aphid species and the two yellow dwarf virus species of greatest concern in Europe to examine how diversity at each level of the vector-virus system impacts disease spread. More broadly, our modelling framework represents a tool that can be used to explore vector-virus-host interactions in greater detail, and we use cereal systems as a case study. Our modelling scenarios highlight diversity at each scale as an important factor. We show that the two main vector species have contrasting dispersal patterns, and that this interacts with virus transmission efficacy to influence disease spread; genetic variation within a vector population is a key driver behind disease spread and disease risk; and for vector species that transmit multiple virus species, the specific vector-virus species combination is a key risk factor to consider.

## Introduction

Cereals (wheat, barley, oats) are some of the most agriculturally important crops grown worldwide, with wheat providing 25% and 23% of the daily total energy intake for the UK and Ireland, respectively (Mottaleb et al., 2022). Across a growing season, cereal crops are exposed to numerous herbivorous pests. These pests have the potential to cause significant crop damage resulting in high yield losses (Nancarrow et al., 2021; Perry et al., 2000; Rogers et al., 2015; Zhao and Zhou, 2024). Of the range of pests that infest cereal crops, cereal aphids are of significant agricultural concern (Leybourne et al., 2024a). On average, cereal aphids can reduce crop yields by 5 *−* 20% through damage caused during aphid feeding (Dedryver et al., 2010). Honeydew secretion by aphids also promotes saprophytic fungi colonisation, reducing the photosynthetic capacity of the plant, having cascading consequences on yield production(Rabbinge et al., 1981).

In Europe, the grain aphid (*Sitobion avenae*), the bird cherry-oat aphid (*Rhopalosiphum padi*), and the rose-grain aphid (*Metapolophium dirhodum*) are the dominant cereal aphid species. These species are vectors of several devastating cereal pathogens, including the yellow dwarf viruses, significant aphid-vectored viruses affecting cereal crops (Aradottir and Crespo-Herrera, 2021; Leybourne, 2024a).

Yellow dwarf viruses are a diverse group of persistent, circulative, non-propagative, phloem-limited viruses (Gray et al., 2014). Prominent yellow dwarf viruses include barley yellow dwarf virus (BYDV, Tombusviridae: Luteovirus), cereal yellow dwarf virus (CYDV, Solemoviridae: Polerovirus), maize yellow dwarf virus (MYDV, Solemoviridae: Polerovirus), and wheat yellow dwarf virus (WYDV, Solemoviridae) (Leybourne, 2024a). Infection with these viruses cause yellow dwarf disease, the main symptoms of which include crop stunting, shrivelled grain, and chlorosis (Leybourne, 2024a). Depending on crop variety, viral pressure, and/or time of infection, yellow dwarf virus-associated yield penalties can be severe, with yield losses of 20 *−* 80% reported (Liu et al., 2014; Nancarrow et al., 2021; Perry et al., 2000).

Primary infection with yellow dwarf virus occurs when virus-carrying winged (alate) aphids migrate into recently sown crops (Leclercq-Le Quillec et al., 2000; Plumb, 1976). These alate aphids subsequently feed and reproduce before moving to new plants or migrating to nearby fields. This behaviour leads to sporadic patches of primary virus infection within the field (Halbert and Pike, 1985). The offspring of these migratory aphids are often wingless (crawling) and develop to adulthood feeding on the virus-infected plants they are born on. As a result, this second generation of aphids have a high likelihood of acquiring the virus and being rendered viruliferous. Secondary (localised) spread of the virus is caused by movement (crawling) of these aphids within the field and across the crop (Halbert and Pike, 1985). The yellow dwarf virus management strategy currently followed by growers in the UK and Ireland is to consider a management intervention if a single cereal aphid is found within the crop before growth stage 31 (Leybourne et al., 2024a) – once the crop reaches growth stage 31, the plant can naturally tolerate yellow dwarf virus infection (Doodson and Saunders, 1970). However, these thresholds have not been tested under field conditions and a low threshold likely contributes to increased application of management interventions such as insecticide treatments. Current guidance is to apply the same yellow dwarf virus threshold to all vector species, all populations within a vector species, and all virus species.

In total, there are around seven described BYDV species, two CYDV species, one MYDV species, one WYDV species, and one species unassigned a genus (Aradottir and Crespo-Herrera, 2021). BYDV-PAV (Tombusviridae: *Luteovirus pavhordei*) and BYDV-MAV (Tombusviridae: *Luteovirus mavhordei*) are considered the most agriculturally impactful yellow dwarf virus specie. The dominance of specific yellow dwarf virus species differs between regions – in mainland Europe BYDV-PAV is thought to be the most abundant, whereas in the United Kingdom BYDV-MAV and BYDV-PAV occur at similar levels (Foster et al., 2004) and in Ireland BYDV-MAV is considered the dominant species (Byrne et al., 2024). However, many of these surveys would have been unable to differentiate between two related BYDV species, BYDV-PAV and BYDV-PAS (Tombusviridae: *Luteovirus pashordei*), potentially leading to an overestimation of BYDV-PAV abundance and an underestimation of BYDV-PAS. Recent advances in molecular diagnostics can more readily discriminate between these related species (Byrne et al., 2024). Therefore, updated epidemiological surveys using testing methods with greater precision are required in order to revise our understanding of species dominance. Nonetheless, yellow dwarf virus incidence is sporadic in nature and the prevalence of species can vary within regions (Dempster and Holmes, 1995; Liu et al., 2020) and fluctuate between monitoring years (Liu et al., 2020).

Each cereal aphid species can transmit a range of yellow dwarf virus species, however the efficiency with which a yellow dwarf virus species is transmitted by each vector species is highly variable (Aradottir and Crespo-Herrera, 2021; Leybourne, 2024a). For example, *R. padi* is the most efficient vector of BYDV-PAV but is broadly inefficient in vectoring BYDV-MAV (Jing-Quan et al., 1997), conversely *S. avenae* is the most efficient vector of BYDV-MAV and is moderately efficient at vectoring BYDV-PAV, while *M. dirhodum* can effectively transmit both BYDV-PAV and BYDV-MAV (Leybourne, 2024a). The efficacy of disease transmission can also vary significantly between discrete populations (e.g., distinctive genotypes) within a given vector species (Leybourne et al., 2024b). Due to emerging resistance and reduced sensitivity to insecticides, cereal aphids are becoming increasingly important agricultural pests (Fontaine et al., 2023; Foster et al., 2014; Walsh et al., 2020). This is increasing the need for more sustainable pest management options (McNamara et al., 2020) and a better understanding of how diversity within aphid populations affects agriculturally important phenotypic traits.

Decision support tools are tools, or systems, that support farmers and growers in making pest and disease management decisions. Most decision support tools developed for aphid and yellow dwarf virus management support growers by optimising the use of management interventions, primarily the timing of insecticide applications (Leybourne et al., 2024a). The majority of decision support tools rely on statistical or mathematical models to estimate various aspects of pest and disease phenology (Leybourne et al., 2023). There are several decision support tools currently available for cereal aphid and yellow dwarf virus management, these include models that: predict aphid activity and virus risk (Kendall et al., 1992; Morgan, 2000); estimate aphid arrival, yellow dwarf virus incidence, and yield loss (Thackray et al., 2009); and estimate secondary virus spread (Leclercq-Le Quillec et al., 2000). However, many of these are limited in scope and focus mainly on one cereal aphid species (primarily *R. padi*) and one virus species (primarily BYDV-PAV) at a time. Although the insights gained by these models are useful, and the decision support tools themselves are seen as an integral component of integrated pest management, there are various nuances within the vector-virus system that are not considered by current decision support tools. Key factors not currently considered include: Variation in disease transmission between different vector populations; how variable life-history traits impact vector dispersal and disease dynamics; insecticide resistance status of the aphid; and natural prevalence of yellow dwarf virus within the local aphid population – currently all cereal-colonising aphid species found within the crop are assumed to carry yellow dwarf virus. If we can integrate these nuances into future decision support tools, then we can develop more sustainable means of pest and disease management.

Bespoke management options based on the composition of local vector-virus populations represent novel avenues that could be explored to develop more sustainable pest and disease management strategies. Different cereal aphid species vary in their capacity to vector each yellow dwarf virus species (Aradottir and Crespo-Herrera, 2021; Leybourne, 2024a) and populations within a species (i.e., clonal populations and genotypes) differ strongly in their agricultural importance, including virus transmission capacity, population growth, insecticide resistance, and production of winged morphs (Leybourne et al., 2024b; Mahieu et al., 2024). However, these factors are not considered in current management strategies (Leybourne et al., 2024a). As these phenotypes are heritable, this knowledge could be used to develop targeted and sustainable management approaches that are centred on the risk presented by the local vector community, instead of the current zero-tolerance approach. Developing models that can predict vector dispersal and yellow dwarf virus spread for discrete populations (i.e., variability between the main species and variation amongst distinctive genotypes within a species) would enable more bespoke management approaches to be followed that are based on the risk presented by the local vector-virus population.

Here, we describe a stage-structured model that can estimate vector dispersal and disease spread. We use data for two of the main yellow dwarf virus vector species, *R. padi* and *S. avenae*, and the two yellow dwarf virus species of greatest concern in Europe, BYDV-MAV and BYDV-PAV, to examine how diversity at each level within the vector-virus system impacts vector dispersal and virus spread. We specifically focus on three levels of diversity, tested through three distinctive modelling scenarios to answer three fundamental questions:

1. How do the main vector species differ in their capacity to transmit the dominant BYDV species, BYDV-PAV, and how does this influence disease dynamics?
2. How does genetic diversity within vector species influence disease transmission, and what are the consequences of this for secondary disease spread?
3. How does variation in vectoring capacity for different virus species within a single vector species impact disease transmission and disease spread?

These modelling approaches are designed to increase our understanding of vector-virus-crop dynamics and allow us to model specific biological scenarios. However, significant knowledge gaps remain that need to be filled before we can fully capture, and model, system interactions and complexity in a comprehensive manner that would support the development of decision support tools. We highlight several of these knowledge gaps as focal areas for further exploratory research, with the ultimate intention of leading to the development of the next generation of decision support tools.

## Materials and Methods

### Data sources for model parameterisation

Our stage-structured model (described below) is parametrised using empirical data collected by our research teams or sourced from the wider literature. BYDV-PAV transmission efficiency for *R. padi* and *S. avenae* was provided by Leybourne (detailed in Leybourne et al. (2024a)); BYDV-MAV transmission efficiency for *S. avenae*, life-history traits for *S. avenae*, and BYDV-MAV prevalence data were provided by Schughart, Mostafiz & McNamara (preliminary data); life-history data for *R. padi* and *S. avenae* was provided by Leybourne (preliminary data); BYDVPAV prevalence data was provided by Bramham (preliminary data); and overall prevalence data of BYDV in *R. padi* and *S. avenae* were obtained from White et al. (2023). Figure one show the infection pathways we model in our system.

### Model Notation

In our model description we refer to winged (alate) adult aphids as “flying adult”, and unwinged (apterous) adult aphids as “crawling adult”. This is to enable simplified model notation, through single character notation of f and c. We also refer to juvenile aphids as either “flying juvenile”or “crawling juvenile” this is to reflect whether the juvenile will develop into a flying or walking adult.

### Model Construction

Estimates of the relative prevalence of aphids carrying MAV and PAV exist from the Rothamsted Research Insect Survey in the UK and Teagasc monitoring in The Republic of Ireland. For the case of *R. padi* transmission efficiency of BYDVPAV, this can vary between aphid genotypes (Leybourne et al., 2024b). However, *R. padi* is inefficient in transmitting BYDV-MAV (Leybourne, 2024a). Conversely, *S. avenae* can transmit both BYDV-MAV and PAV with variable efficacy. We therefore develop a two species (MAV and PAV) model for *S. avenae* and a single species (PAV) model for *R. padi*. We consider an explicitly spatial model where a field is made up of cells and the population within each cell is modelled using a stage structured model with two aphid phenotypes crawling (wingless) which can disperse only by dropping off a plant and crawling, and flying which can fly between plants and an explicit juvenile stage where juveniles carrying viral species *k* = *p, m* (PAV or MAV carrying) of phenotoype *i* = *c, f* (crawling of flying) are represented by *J_k_* and adults by *A_k_*. Following the Methodology of Gurney and Nisbet (1998) we define the population change in each cell as follows

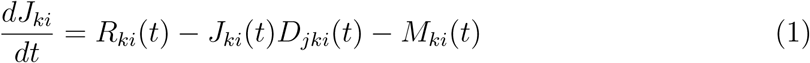

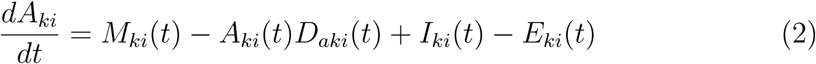

Where *R_ki_* represents recruitment (or reproduction) to phenotype *k* of species *i*, *D_ki_*, *M_ki_* maturity, *I_ki_* immigration to a patch and *E_ki_* emmigration from a patch. We will now describe each component of the model in turn

### Recruitment

We assume that aphid reproduction is logistic with a maximum reproductive rate *r* and carrying capacity *C* and that natural predators remain at a level sufficient to maintain a per capita rate of aphid consumption, *δ*. Adult flying phenotypes are assumed to produce only crawling juveniles whereas crawling adult phenotypes are assumed to produce both flying and crawling phenotypes with the relative proportions dependent on aphid density. Crawling morphs produce flying morphs with a density dependent probability 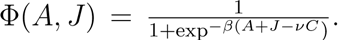. This produces a sigmoid curve running between 0 and 1 where *ν* is the proportion of carrying capacity at which there is a 50% chance of producing winged morphs and *β* controls the steepness of the curve. See figure 2 for examples of what this means. Juvenile and adult aphids of all phenotypes utilise the same resources so density dependent reproduction is a function of both nymphs and adults of all phenotypes. Thus, if *A* is the total number of adults of all species, and *J* the total number of juveniles, the equation for reproduction of crawling morphs of genotype *k* is

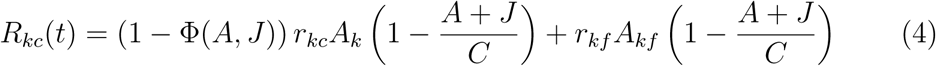

**Figure 1:**
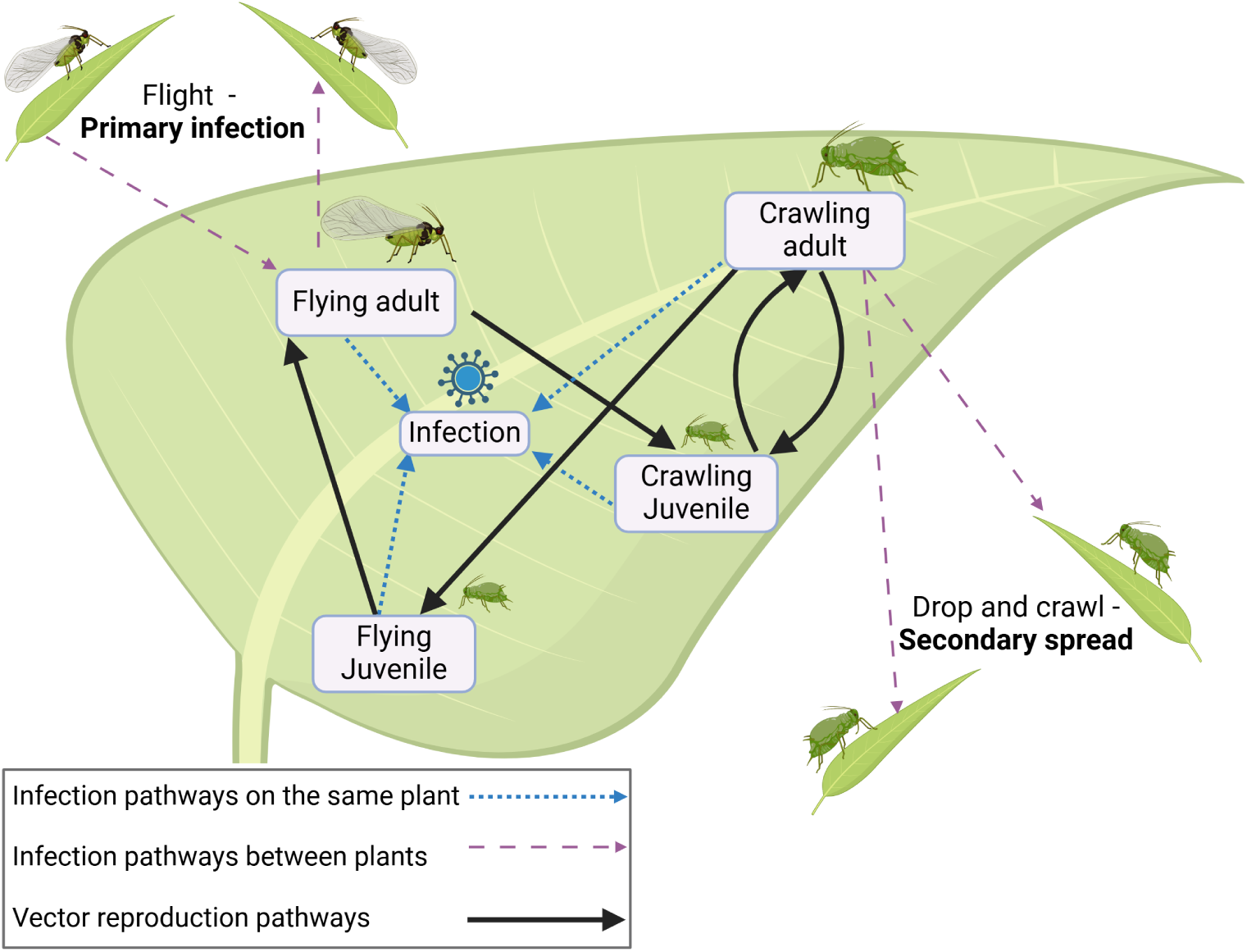
Graphical overview of the model infection pathways.

**Figure 2:**
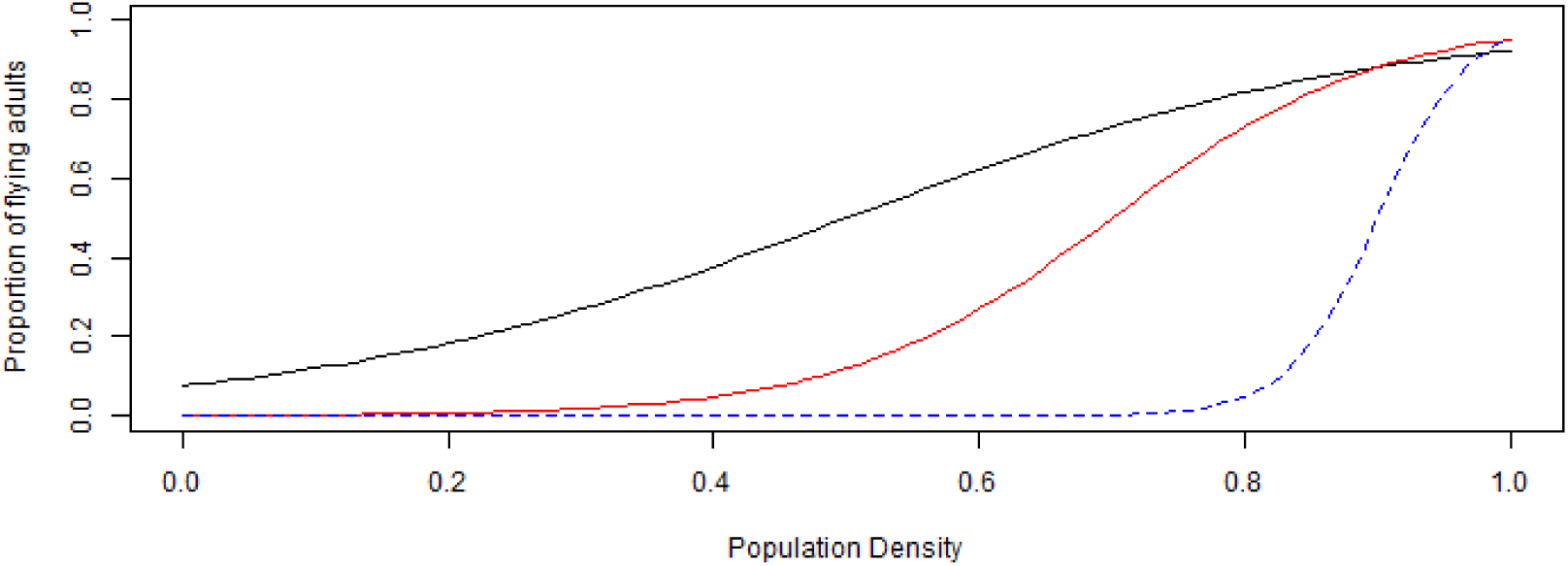
Relationship between flying-morph production and aphid density. The solid black line shows the values for *R. padi*, genotype 1 & 2 (*β* = 5*, ν* = 0.5). The dotted red line shows the values for *R. padi* genotype 3 (*β* = 10*, ν* = 0.7).The blue dashed line shows the values for *S. avenae* (*β* = 30*, ν* = 0.9).

and reproduction flying morphs is defined by

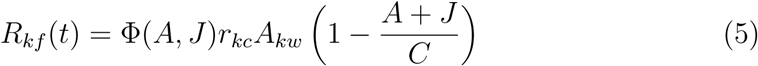

where *r_kc_*and *r_kf_* are the the intrinsic reproductive rates of genotype *k* crawling and walking morphs respectively.

### Maturation and Death

Juveniles of phenotype *i* have a development time *τ_i_*. A constant background mortality rate of *δ_n_* is assumed with an additional mortality rate *δ_h_* for adults is assumed to be independent of whether the aphid carries the yellow dwarf virus. so

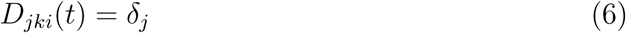

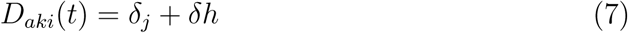

Juveniles which do not die mature as adults after time *τ_i_* so we have

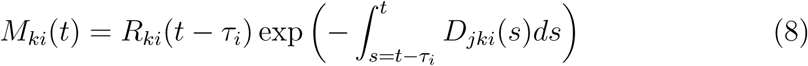

### Natural virus infection

White et al. (2023) found that proportions of aphids carrying BYDV in suction trap captures was consistent with in-field yellow trap captures, with an average BYDV prevalence of 20%. It is therefore reasonable to assume that aphid populations are well-mixed with respect to BYDV and to model transmission rates as a fixed proportion of aphid population density in a given cell *ρ* multiplied by a vector species or genotype-defined transmission efficiency *γ*. We assume that there are a fixed number of plants per cell (*cellsize*^2^) and that there is no plant death. We monitor infection in the plants as a simple infection rate subject to the proportion of uninfected plants in the patch and incorporate no recovery. Thus, if *η_k_* is the infection rate of an aphid with vector transmission efficiency *k*,the probability that an uninfected plant escapes infection is

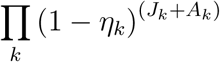

Let *γ_k_*= *log* (11 *− η_k_*) then the probability of an uninfected plant being infected is

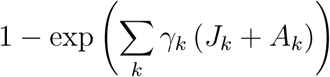

Therefore, the change in infected plants, *P* will be

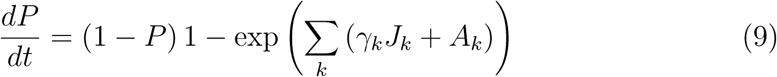

### Immigration and Emmigration

We assume that only adults move and this is monitored by counting the number of aphids emmigrating from a cell and adding up the number of aphids immigrating to that cell from all other cells in the field. Emmigration is modelled as a fixed probability *µ_i_*, *i* = *c, f* of crawling or flying between plants with a dispersal kernel following a truncated normal distribution with dispersal kernels *σ_c_* and *σ_f_* and truncations of *m_c_*, and *m_f_* for crawling and flying adults respectively. Where the *m_i_* is the maximum number of plants that an aphid of phenotype *i* can travel can travel. Thus the dispersal kernel and truncation will be 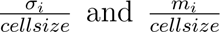respectively.

Immigration to a given cell is the sum over all cells of emmigration to that cell multiplied with a fixed probability *α_i_*, *i* = *c, f* of successfully crawling and flying phenotypes finding a plant in that cell on arrival.

To summarise we have

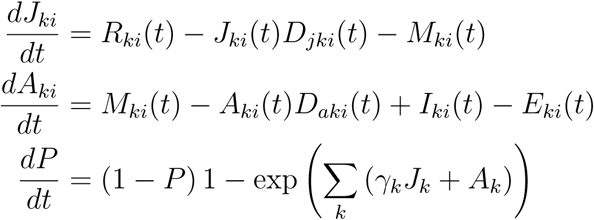

with

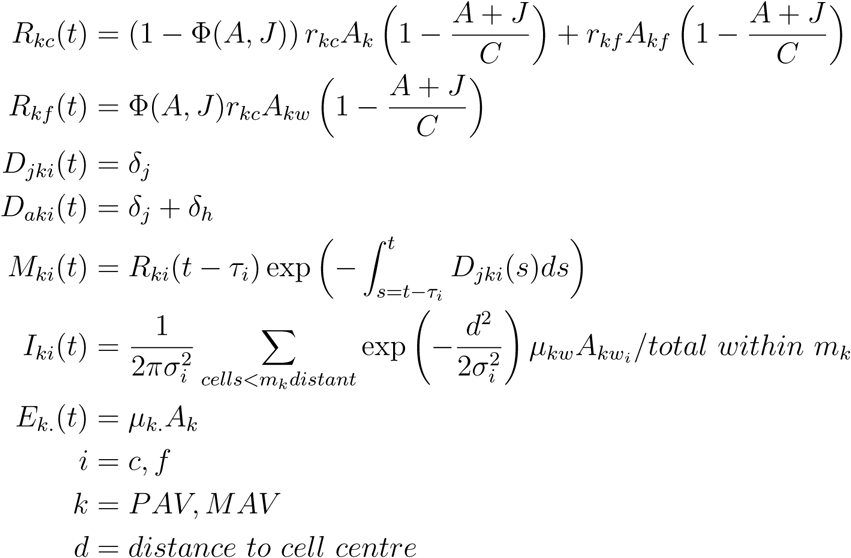

### Parameterisation

The advantage of the model as formulated is that many of the raw parameters are obtainable from available empirical investigations. The key parameters for the model are summarised in table 1 and their derivation is discussed below.

**Table 1:** Model parameter values.

| parameter | description | *R. padi*<br>PAV | *S. avenae*<br>PAV | *S. avenae*<br>MAV |
| --- | --- | --- | --- | --- |
| $exp(r_c(r_f))$ | Daily reproductive rate | 3 (2.7) | 9 (8.1) | 9 (8.1) |
| $\delta_n$ | background mortality rate | | 0.018 | |
| $\delta_m$ | additional adult mortality rate | | 0.087 | |
| $\tau$ | juvenile development time | 7 (9) | 9 (10) | 9 (10) |
| $\mu_c$ | P(crawling adult drops) | 0.05/40 | 0.5/40 | |
| $f_c$ | P(crawling aphid finds new plant) | | 0.1 | |
| $f_f$ | P(flying aphid finds new plant) | | 0.8 | |
| $\beta$ | winged morph birth density response | 5, 5, 10 | 30 | 30 |
| $\nu$ | density at which 50% are winged | 0.5, 0.5, 0.7 | 0.9 | 0.9 |
| $exp(\eta)$ | daily BYDV transmission rate | 0.83/30, 63/30, 0.44/30 | 0.16/30 | 0.3/30 |
| $\rho$ | proportion of BYDV carriers | | 0.2 | |
| $\mu_c$ | P(crawling adult drops) | | 0.5/40 | |
| $\mu_f$ | P(winged adult flies) | | 1/40 | |
| $\sigma_c$ | dispersal kernel of crawling adults | | 5 | |
| $\sigma_f$ | dispersal kernel of flying adults | | 50 | |
| $\alpha_c$ | probability that a dropped aphid successfully migrates | | 0.5 | |
| $\alpha_f$ | probability that a flying aphid successfully migrates | | 0.8 | |
| $m_c$ | max plant dispersal of crawling adults | | 25 | |
| $m_f$ | max plant dispersal of flying adults | | 250 | |
| $cellsize$ | no. plants per cell side | | 25 | |
| $runtime$ | days for which the model runs | | 90 | |

### Adult reproduction and flying morph production

Over the course of their life it can be assumed that crawling *R. padi* adults produce an average of 3 nymphs per day, with a slightly lower rate of 2.7 nymphs per day for flying phenotypes (calculated from open-access data available at Leybourne (2024b)). This equates to an instantaneous reproductive rate of *r_c_*= ln(3) and *r_f_* = ln(2.7). Similarly flying and crawling *S. avenae* adults can be assumed to produce a greater number of nymphs per day, on average 9 and 8.1 nymphs, equating to instantaneous reproductive rates of *r_c_* = ln(9) and *r_f_* = ln(8.1). With regards to the production of flying adult morphs, on average *R. padi* produce more winged morphs when compared with *S. avenae* Leybourne (2024b). We also assume that there is a density-dependent increase in flying morph production so in *R. padi*, with flying morphs produced even at quite low densities with a steady increase as population density increases. For *S. avenae*, we assume littleto-no flying morph production until very high populations densities when there is a sudden switch to almost exclusive flying morph reproduction (A Karley *pers. comm*). We therefore set the density at which 50% of morphs are winged to be 50% of maximum capacity, *ν* = 0.5, for *R. padi* and 90% of maximum capacity, *ν* = 0.9 for *S. avenae* with *β* = 0.5 creating a gradual increase in density for *R. padi* and *β* = 30 creating a very steep increase in winged morphs for *S. avenae* (Figure 2).

### Maturation and Death

Estimates of juvenile development *τ_.k_* are taken from observations on the prereproductive development time for *R. padi* Leybourne (2024b) with observations for *S. avenae* comprising preliminary data (Mostafiz, Mc Namara) .

There is no field data for *R. padi* mortality and very limited data for other species of aphid. However, following the example in Preedy et al. (2020) we use a combination of survival data from laboratory and caged-field experiments to estimate daily mortality rates for juveniles and adults.

Leybourne et al. (2018) found little-to-no *R.padi* nymph mortality after 7 days in glasshouse conditions so we use the field data to estimate background natural mortality for all aphids. Watt (1979) and Howard and Dixon (1992) found values of 87 *−* 95% survival of cereal aphids on immature plants and 25 *−* 40% on mature plants. Based on these data we assume that, in the field approximately 85% of juveniles *R. padi* survive to reproductive maturity after 9 days which implies a background natural mortality rate *δ_j_* = 0.018 per nymph per day. Aqueel and Leather (2011) found total longevity of *R.padi* to be between 28.1 and 30.8 days with a development time of between 5.6 and 6.4 days in controlled environment conditions of 22 deg *C* with a 16:8 day/night regime and only one aphid nymph per plant. This suggests adult survival of 22-24 days under optimum conditions. Leybourne et al. (2018) found average adult *R. padi* survivorship after 21 days of 10 *−* 20% in glasshouse conditions with a development time of between 7 and 9 days. Aqueel and Leather (2011) also consider *S. avenae* finding development times 7.2-7.6 days and total longevity of between 34.4 and 36.3 days suggesting an adult survival time of 27-29 days under optimal controlled environment conditions. Field conditions where aphids are competing for plant resources and are subject to fluctuating environmental conditions are likely to lead to decreased longevity and we note the longer development times in the data set at Leybourne (2024b) would make the adult lifespan of *S. avaenae* similar to that of *R. padi*. We therefore assume the same adult lifespan for both species, calculating mortality rates as follows. It is reasonable to assume that the same background field mortality that impacts juveniles will also apply to adults. The adult death rate of aphids (*δ_m_*) is therefore calculated from 2 components; a base rate of 0.087 per day so that 14.7% of individuals are alive after 21 days alongside a background field mortality rate of 0.018, calculated from juvenile survival. Thus *δ_h_* = 0.087 + 0.018 = 0.105 so that after 21 days 9.7% of individuals are alive.

### Virus transmission efficiency and daily transmission rate

Virus transmission efficiency for *R. padi* and *S. avenae* when vectoring BYDVPAV are taken from Leybourne et al. (2024b), with transmission efficiency for *S. avenae* when vectoring BYDV-MAV taken from preliminary data (Schughart, Mc Namara). On average the longevity of aphids is approximately 30 days so the daily transmission rate from the aphid to a a susceptible plant, *η* is virus transmission efficiency divided by 30 and the instantaneous rate log of that value.

### Immigration and Emmigration

These parameters have not been directly measured, but have been selected as being reasonable based on observations of aphid behaviour during fieldwork focussed on other behavioral traits. We assume different tendencies to drop from plants with *µ_c_* for *R. padi* at 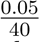 representing a tendency for approximately 5% of wingless adults to drop from plants whereas 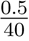 represents represents observations that as many as 50% of wingless *S. avenae* adults can be found on the ground over the course of a season.

We assume that all winged adults will fly with daily probability flight being 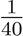 for flying adults from both species. In the absence of species-specific aphid dispersal measurement, we assume the same dispersal kernels for crawling and flying adults. It is flying will successfully alight on a new plant with probability *α_f_*= 0.8 and that adult crawling morphs successfully finding crawling onto another plant with probability *α_c_* = 0.1. The maximum dispersal of crawling morphs *m_c_* is 25 plants and of flying morphs *m_f_*is 250 plants.

### Model Structure

The model is run for 90 days the time during which most cereals are vulnerable to infection following crop sowing (Leybourne et al., 2024a). We use a field of 100 *×* 100 square cells with a side length of 25 plants. This equates to 625 plants per cell and at a planting density of 150 plants per *m*^2^ a cell length is approximately 2 meters allowing for realistic representation of a 200 m by 200m field.

## 1 Results

### Variable life-history strategies between two vector species determine vector dispersal and disease spread

In our modelling scenarios we see a more rapid population increase for *S. avenae* compared with *R. padi*, with *R. padi* also producing a greater proportion of winged aphids as the season progresses (Figure 3). This inter-species variation in phenology and demography produces markedly different vector dispersal patterns for the two species (Figure 4). For *S. avenae* we observe a clear and pronounced moving front of crawling aphids, whereas for *R. padi* we observe thinly spread, but densely clustered, populations of crawling aphids with colony dispersal primarily through within-field migration of the winged aphids (Figure 4). Dispersal at 45 and 60 days is displayed in the supplementary material Figure S1 and Figure S2, respectively.

**Figure 3:**
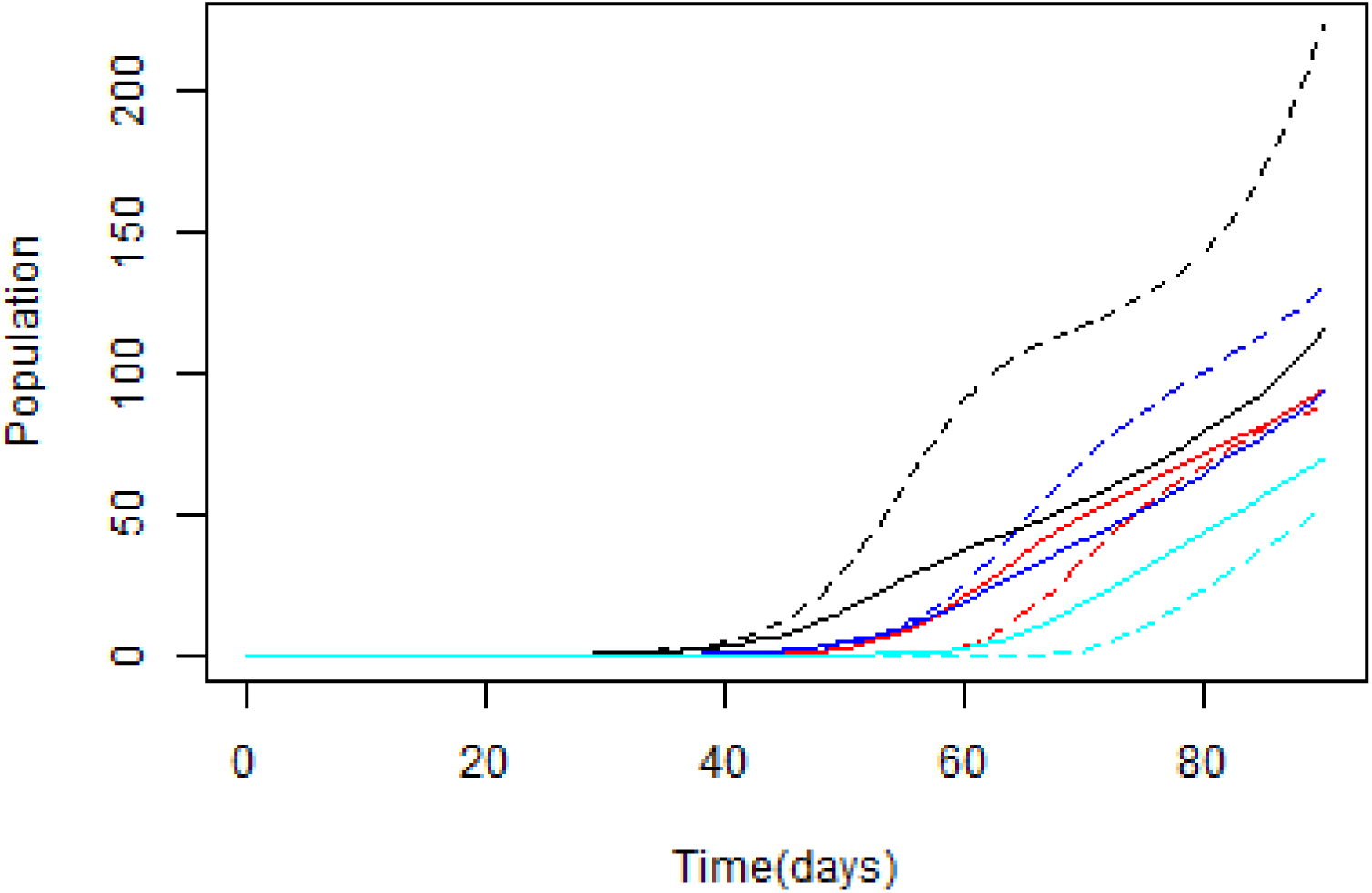
Model outputs showing estimations of population change over the 90-day season. Solid lines represent *R. padi* and dashed lines represent *S. avenae*. Black lines represent crawling juveniles; red lines flying juveniles; blue lines crawling adults; and cyan lines flying adults.

**Figure 4:**
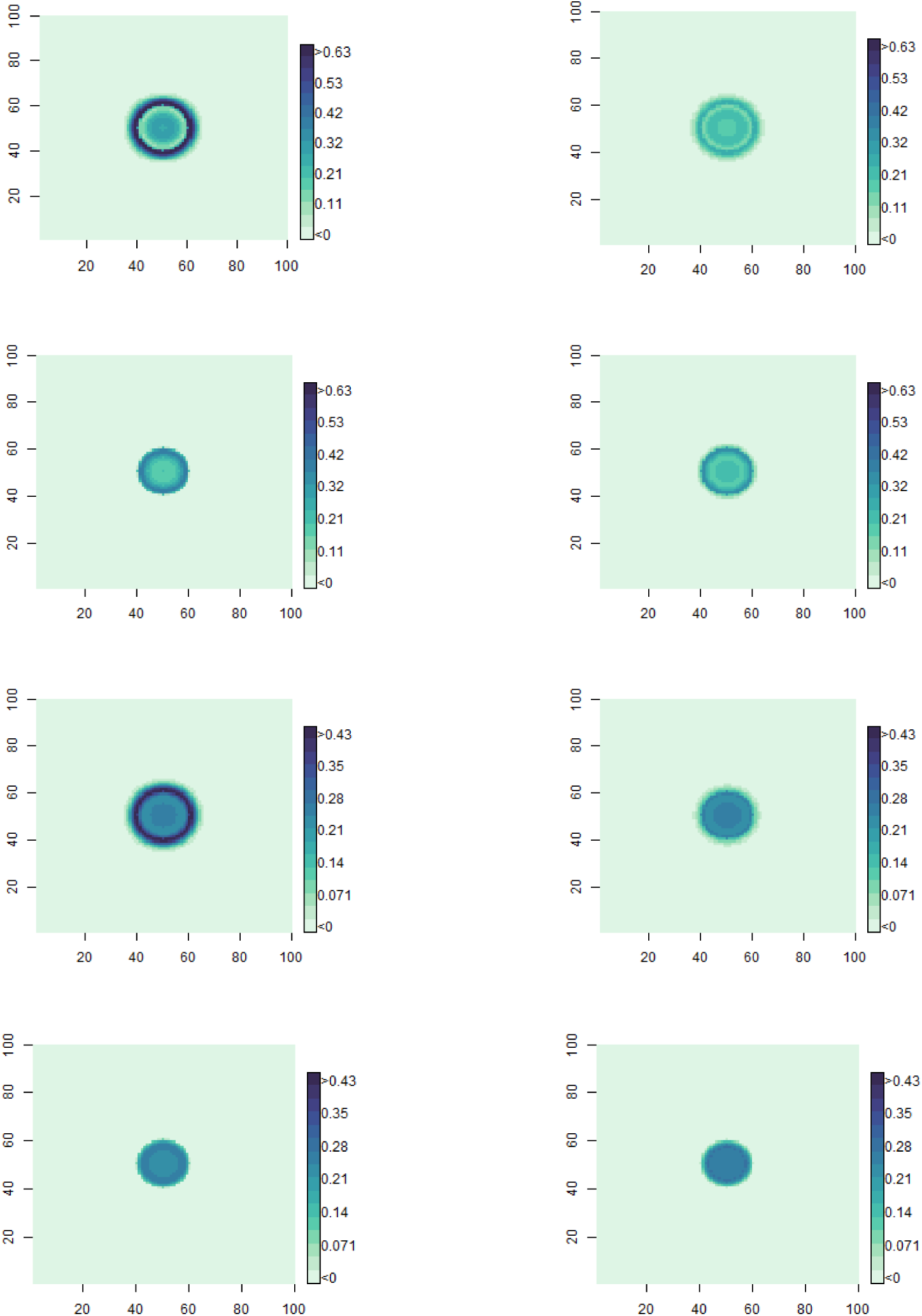
Model outputs showing estimations of vector dispersal after 90 days. Left column shows *S. avenae* and right column *R. padi*. Top to bottom: Crawling juveniles, flying juveniles, crawling adults, flying adults. Juveniles share the same scale [0, 0.632] and adults share the same scale [0, 0.425].

This inter-species variation in vector demography (Figure 3) and dispersal (Figure 4) also produces contrasting patterns of population distribution and BYDVPAV disease incidence as the distance from the primary inoculation point increases over the modelled 90-day period (Figure 5). This results in contrasting disease spread: In the *S. avenae* model we observe comparatively little disease infection over the 90 days when compared with *R. padi* (Figure 6). Although *S. avenae* populations increase at a greater rate, the lower capacity to transmit BYDV-PAV results in a significantly lower disease risk when compared with *R. padi*. For *R. padi*, we observe patterns of disease incidence that broadly follow the vector dispersal pattern (Figure 6). The profile of population and infection densities at distance from the initial inoculation is displayed in Figure 5 highlighting demographic variation between the aphid species, with *S. avenae* represented by consistent movement of crawling aphids and a steady increase in population density.

**Figure 5:**
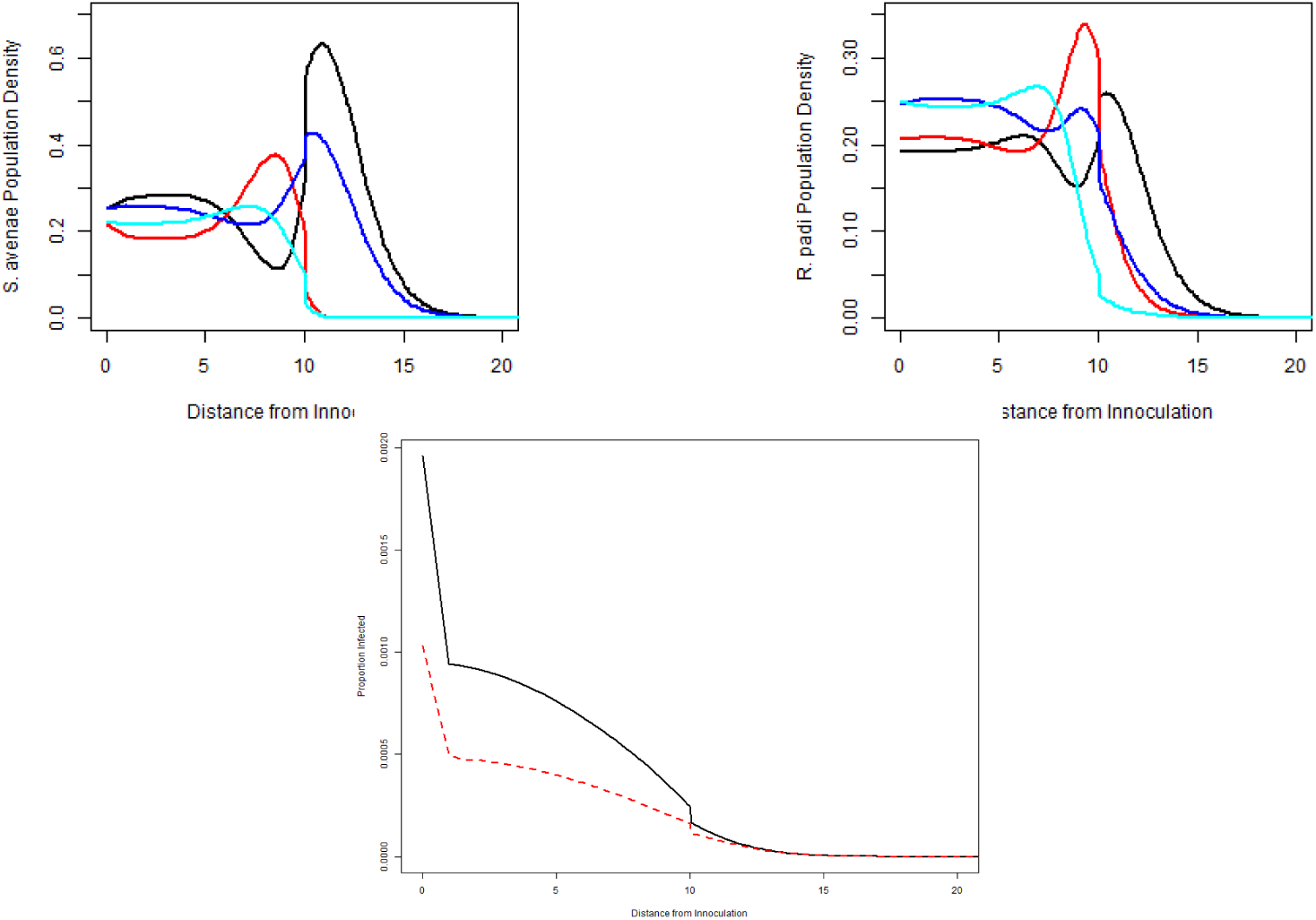
The top figures show population density at distance from the initial inoculation after 90 days. The left figure shows *S. avenae* and the right figure *R. padi*. Crawling juveniles are represented by black lines, flying juveniles red lines, crawling adults blue lines and flying adults cyan lines. Note that the density scale for *R. padi* is half that of *S. avenae*. The bottom figure shows infection at distance from the initial inoculation after 90 days. The solid line shows infection from *R. padi* and the dashed line infection from *S. avenae*.

**Figure 6:**
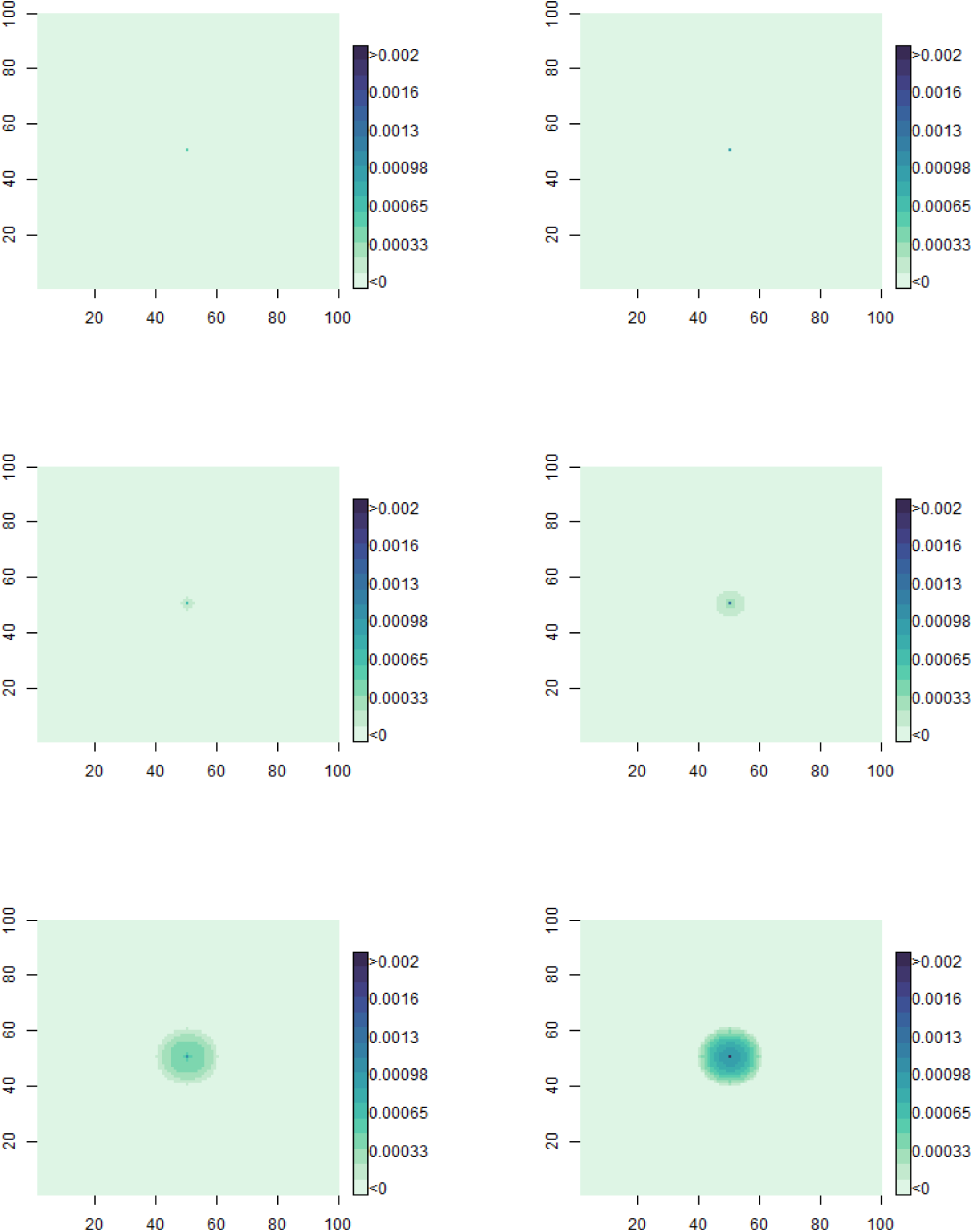
Model outputs showing estimations of BYDV-PAV disease infection for the two vector species across 90 days. Left column shows *S. avenae* and right column *R. padi*. Top to bottom, disease infection after: 45 days, 60 days, 90 days.

### Heritable phenotypes within an aphid species produce different disease infection scenarios

Using data showing variable virus transmission phenotypes between *R. padi* genotypes we modelled the effect intra-species variation has on BYDV-PAV infection. Our model shows that this variable transmission phenotype has a significant impact on total disease incidence within the field (Figure 7), with the genotypes that transmit BYDV-PAV with higher efficiency infecting a greater number of plants when compared with the other genotypes. When compared with *S. avenae* transmission, all *R. padi* genotypes infected a greater proportion of plants with BYDV-PAV than *S. avenae*. However, only one *S. avenae* genotype was tested.

**Figure 7:**
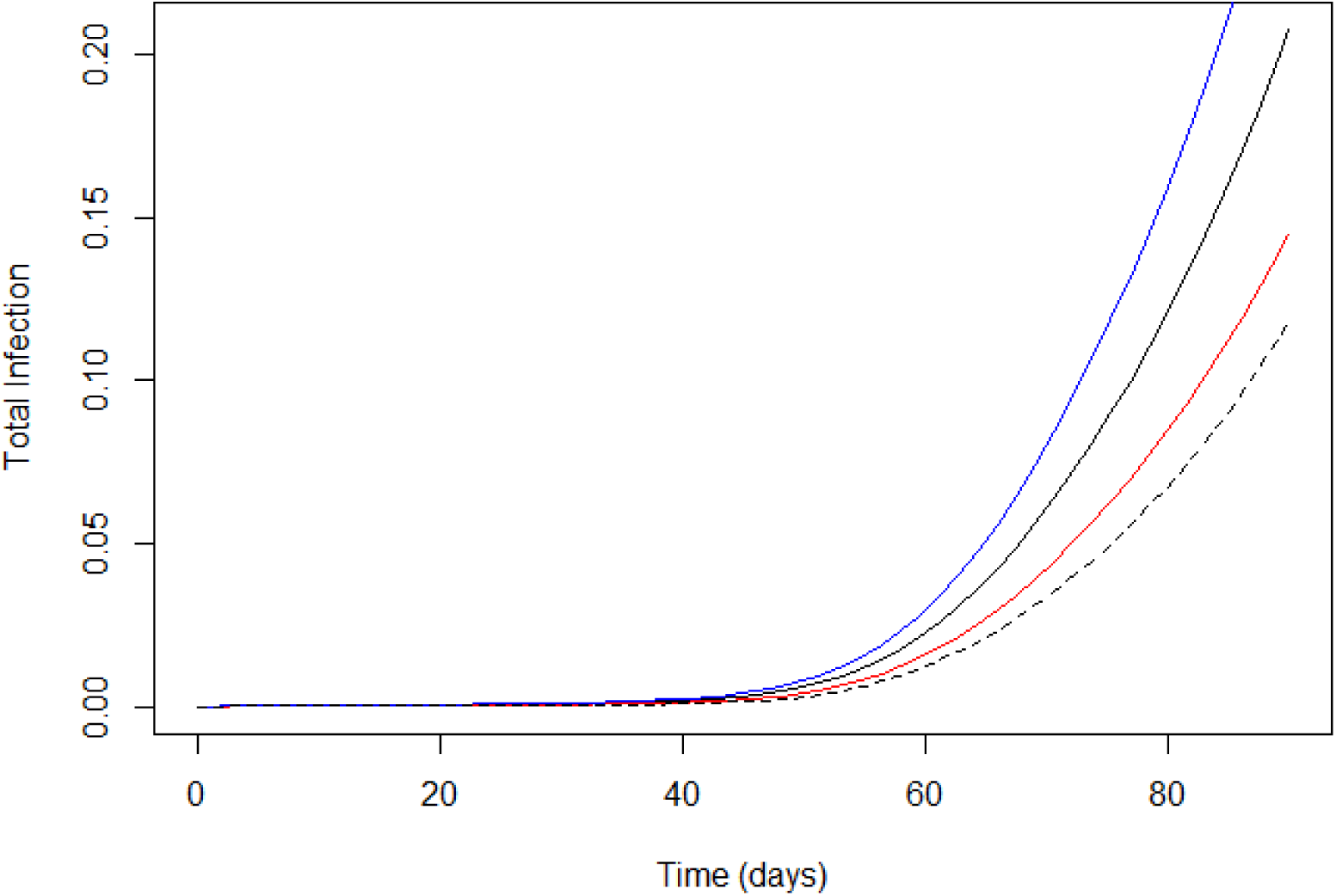
Model outputs showing estimations of disease infection for three *R. padi* genotypes with variable BYDV-PAV transmission probabilities (blue = high transmission [0.83 probability], black = moderate transmission [0.63 probability], red = low transmission [0.44 probability]); dashed line represents *S. avenae* for comparison.

### Virus-vector species combinations are an important driver of disease success

Using data reporting variable virus transmission of BYDV-MAV and BYDV-PAV for *S. avenae* (high transmission efficiency of BYDV-MAV, low transmission efficiency of BYDV-PAV), we modelled the effect these vector-virus combinations have on disease infection. Our model shows that disease progression and spread varies between the two virus species (Figure 8), with BYDV-MAV spreading more strongly than BYDV-PAV.

**Figure 8:**
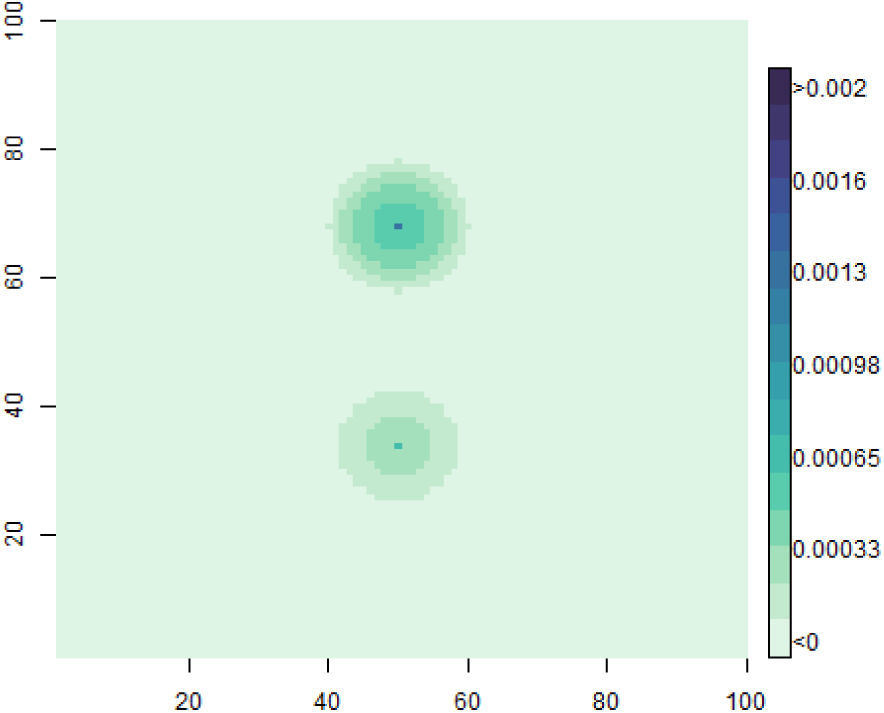
Plant infections from the two different species of BYDV in *S. avenae* 90 days after innoculation. The top innoculation is from MAV carrying aphids and the bottom from PAV carrying aphid.

## Discussion

We have developed a modelling framework structured to allow the inclusion of multi-trophic, multi-species, multi-stage interactions in discrete-space and continuous time allowing additional aspects of system complexity to be incorporated or removed as needed. Model code is freely available at https://gitlab.bioss.ac.uk/natural-pest-management/BYDVmodel. We use this framework to model vector-virus-host interactions and examine how three levels of diversity (vector inter-species diversity, vector intra-specific diversity, and virus diversity) impacts vector dispersal and disease spread in an agriculturally important pathosystem. We use the insights generated by the model to answer three fundamental questions:

1. How do the main vector species differ in their capacity to transmit the dominant BYDV species, BYDV-PAV, and how does this influence disease dynamics?
2. How does genetic diversity within vector species influence disease transmission, and what are the consequences of this for secondary disease spread?
3. How does variation in vectoring capacity for different virus species within a single vector species impact disease transmission and disease spread?

Modelling vector and disease dispersal is a fundamental principle of integrated pest management, and several models have been developed to support cereal aphid and yellow dwarf virus management. The majority of these focus on predicting aphid abundance, arrival, or migration, and use these to estimate subsequent yellow dwarf virus risk (Harrington et al., 1991; Kendall et al., 1992; Lankin-Vega et al., 2008; Morgan, 2000; Thackray et al., 2009). Models that predict secondary spread of several yellow dwarf virus species (BYDV-PAV, BYDV-MAV, and CYDV-RPV) have also been developed (Leclercq-Le Quillec et al., 2000). However, most models focus on modelling interactions, and/or crop risk, for single vector-virus combinations (primarily *R. padi* and BYDV-PAV). This likely leads to an overestimation of vector and disease risk as the different aphid species have variable life-history traits: Generally, when compared with *S. avenae*, *R. padi* are faster to develop and produce a higher proportion of alate aphids. These variable traits are not currently considered when determining yellow dwarf virus risk, and a blanket approach is often applied to all cereal aphid species with advice to apply insecticide if an individual cereal-colonising aphid is found within the crop (Leybourne et al., 2024a). In Europe, the main cereal aphids are *S. avenae*, *R. padi*, and *M. dirhodum*, and the main yellow dwarf virus species are BYDV-PAV and BYDV-MAV. Here we focussed on *S. avenae* and *R. padi*, as these are the two vectors that are thought to represent the greatest yellow dwarf virus risk to winter-sown cereal crops (Leybourne et al., 2024a) and can effectively transmit BYDV-PAV (Aradottir and Crespo-Herrera, 2021). It should be noted that *M. dirhodum* represents an important yellow dwarf virus vector of spring-sown crops in the UK and Ireland. *M. dirhodum* is an understudied species, and as such there is insufficient phenotypic data to robustly parameterise the model for *M. dirhodum*. However, our modelling framework is stage-structured, dynamic, and adaptable *M. dirhodum*-yellow dwarf virus interactions can be modelled as data become available.

Our modelling indicates that vector species is an important factor underpinning disease transmission, particularly for yellow dwarf virus species that can be transmitted by several vector species with variable competence (e.g., BYDV-PAV). The insights generated by our first modelling scenario (modelling vector dispersal and disease spread for *R. padi*, *S. avenae*, and BYDV-PAV) show that the different life-history traits of the two vector species lead to contrasting spatial dispersal between the two species, with a greater dispersal of crawling adults for *S. avenae*. This is likely driven by greater reproductive output and a higher rate of dropping in *S. avenae*, leading to increased local migration. Our modelling also suggests that the disease risk for BYDV-PAV presented by *S. avenae* is significantly lower than the risk presented by *R. padi*, this is despite *S. avenae* having more pronounced dispersal of crawling (crawling) aphids compared with more clustered and dense populations of *R. padi*; this was also indicated in our models as the proportion of plants infected as the distance from the initial inoculation was lower for the *S. avenae* BYDV-PAV model than the *R. padi* BYDV-PAV model, despite *S. avenae* having a much higher overall aphid density . This is likely driven by the much lower vectoring competency of BYDV-PAV by *S. avenae* compared with *R. padi* (Jing-Quan et al., 1997; Leybourne et al., 2024b), and indicates that contrasting disease incidence and spread is primarily driven by variability in disease transmission between the two vector species, and not necessarily secondary dispersal of aphids within the field which might otherwise be considered a primary factor underpinning disease risk (Halbert and Pike, 1985).

Currently, best practice guidelines are to consider a management intervention (e.g., pesticide application) if a single cereal-colonising aphid is found within the crop canopy during the yellow dwarf virus risk period (Leybourne et al., 2024a). However, this is a simplification of the underlying biology as distinct aphid populations exhibit unique agricultural phenotypes, including important traits such as alate production, reproduction, and virus transmission (Gonzalez-Gonzalez et al., 2024; Leybourne et al., 2024b; Mahieu et al., 2024; Ramírez-Cáceres et al., 2019). Many of these phenotypic traits are associated with heritable factors (genotypes and the presence of endosymbiotic bacteria) that can be passed down between aphid generations. With regards to yellow dwarf virus risk, studies have shown that BYDV-PAV transmission efficiency in *R. padi* differs between aphid genotypes (Leybourne et al., 2024b) and observations in the cereal aphid *Schizaphis graminum* found a similar genetic-link underpinning variation in transmission of two yellow dwarf virus species (Burrows et al., 2006, 2007). More generally, there is wide variation in yellow dwarf virus transmission efficiency between aphid clonal populations for multiple vector-virus combinations (reviewed in Leybourne (2024a)). Taken together, these suggest that the disease risk potentially differs between aphid populations, although this hasn’t been empirically tested at the field-scale.

By using phenotype data (development, alate production, reproduction, BYDVPAV transmission) for three *R. padi* genotypes we modelled how phenotypic diversity impacts disease incidence and spread of BYDV-PAV. Variation in BYDVPAV competency in *R. padi* has been well documented (Bencharki et al., 2000; Du et al., 2007; Kern et al., 2022; Rochow and Eastop, 1966), although the interactions between transmission efficacy and other aphid phenotypes that potentially influence disease incidence and spread has not been examined in detail. By linking these phenotype data on virus transmission and aphid life-history in characterised aphid populations with contrasting BYDV-PAV transmission phenotypes, we are able to model these interactions and consequences for disease spread directly. Our modelling indicates that genotype-linked variability in BYDV-PAV transmission efficiency leads to different plant infection rates after the 90-day simulation, with the total number of infected plants almost two-fold higher for the more efficient genotype compared with the least efficient *R. padi* genotype, indicating that variation within discrete vector species is an important factor influencing crop-disease dynamics.

Although we focus on genetic variation and show that the associated phenotypes influence disease incidence and spread, our modelling framework can easily be refined and adapted to include other phenotypes or traits conferred by additional, or iterative levels, of intra-species diversity. Of these, aphid endosymbionts represent an important aspect worthy of future consideration. Key endosymbiont-conferred phenotypes include increased resistance to natural enemies such as parasitoids wasps (Zytynska et al., 2021), which can have potential consequences on the efficacy of biological control interventions. Aphid endosymbionts have also been described to influence vector-virus relationships, including the transmission of pea enation mosaic virus (Sanches et al., 2023). It is currently unclear whether they influence yellow dwarf virus transmission. Our model framework can be adapted to incorporate this additional level of diversity when data become available.

The yellow dwarf virus species complex comprises several related viruses, and each virus species is transmitted by several vector species with varying efficiency depending on the vector-virus combination (reviewed in Leybourne (2024a)). In our final model we used *S. avenae* and two virus species, BYDV-PAV and BYDVMAV, to examine how compatibility between vector species and virus species impacts disease spread in vectors with the same life-history traits. We chose to focus on these vector-virus combinations as *S. avenae* is a vector species that is present throughout the growing season and is a competent vector (i.e., is capable of transmitting the disease, even with low efficiency) for both BYDV-PAV and BYDVMAV (Jing-Quan et al., 1997; Leybourne, 2024a; Leybourne et al., 2024a), which are broadly considered to be the two virus species of greatest abundance and importance in the UK and Europe (Byrne et al., 2024; Foster et al., 2004). Similar to our observations for the contrasting *R. padi* genotypes, our third modelling scenario indicates that the vector-virus combination is an important factor influencing disease transmission, spread, and risk with our model showing higher disease spread of BYDV-MAV compared with BYDV-PAV. This observation is in-line with variability in vector competency for *S. avenae* when transmitting BYDV-MAV, highly efficient vector (Jing-Quan et al., 1997; Leybourne, 2024a), and BYDV-PAV, low-to-moderately efficient vector (Jing-Quan et al., 1997; Leybourne et al., 2024b), and highlights the importance of considering the vector and virus species populations when estimating disease risk. For example, *S. avenae* carrying BYDV-MAV would represent a higher disease risk than *S. avenae* carrying BYDV-PAV.

### Filling knowledge gaps to support the development of future modelling tools

Interactions between aphid vectors, yellow dwarf virus, and host crops are highly complex, particularly when different vector-virus combinations can underpin vector competency, and diversity between and within vector species can have cascading effects on virus transmission and disease risk for a given virus species. Here, we used three biological scenarios to model how diversity within this system impacts vector and disease dynamics, focussing on vector inter-species diversity (*R. padi* vs. *S. avenae*), vector intra-species diversity (three *R. padi* genotypes), and virus species diversity (transmission of BYDV-PAV and BYDV-MAV by *S. avenae*). Generally, our modelling scenarios highlight the importance of this diversity at each scale as important factors influencing vector dispersal and disease spread.

There are still many aspects of diversity in the immediate vector-virus-host system, and the agro-ecosystem more broadly, that will influence vector dispersal and disease spread; however, these cannot currently be incorporated into modelling scenarios. This is due to lack of empirical data. One key missing link is the consideration of co-occurring yellow dwarf virus species within an individual vector, as this can impact disease transmission, including transmission competency for otherwise inefficient vector-virus combinations (e.g., transmission of BYDV-MAV by *R. padi*can be achieved via co-transmission with BYDV-PAV (QUILLEC et al., 1995)). Despite these knowledge gaps, our stage-structured model represents a starting point and provides a framework from which we can add additional levels of complexity iteratively and explore the impacts of these in the future once more data become available. We highlight several important knowledge gaps below:

- More consideration of the contrasting life-history traits between virus-carrying and non-virus-carrying vectors, and a better understanding of how this impacts vector dispersal;
- Greater understanding of bottom-up effects and the impact crop species (wheat, barley, oats) and cultivar has on vector dispersal and disease transmission, particularly with regards to the use of yellow dwarf virus tolerant germplasm;
- Better characterisation of vector competency and transmission efficiency for understudied vectors (e.g., *M. dirhodum*) and lesser studied yellow dwarf virus species (e.g., CYDV-RPV, BYDV-PAS);
- The potential prevalence of multiple co-occurring yellow dwarf virus species within a single aphid and the impact of this on vector dispersal, disease (co)-transmission, and virus competition;
- More consideration of the influence the wider agro-ecosystem will have on vector dispersal and disease transmission, in particular the impact of natural enemies.

Considering these aspects in greater detail, and gathering the information needed to fill these knowledge gaps, would generate more detailed insights into the BYDV vector-virus-host system and allow exploration of field and landscape scale management interventions under evolving climatic conditions. Integrating these nuances into future decision support tools would support the development of more sustainable pest and disease management strategies.

## Funding acknowledgments

The authors gratefully acknowledge financial Support from the Scottish Government Rural Environmental and Scientific Advisory Service Strategic Research programme for KFP, funding from The Royal Commission for the Exhibition of 1851 through a Research Fellowship to DJL (RF-2022-100004), funding from the European Union’s Horizon 2020 research and innovation programme (Marie Sklodowska-Curie grant agreement no.101106698 — MONET) to MMM, and Taighde Éireann, Research Ireland (grant number 22/FFP-A/11049) to MS and LMN. They thank University of Liverpool, Institute of Infection Veterinary and Ecological Science for supporting the project by providing workshop funding through the Network Activity Fund.

## References

1. MA Aqueel and SR Leather. Effect of nitrogen fertilizer on the growth and survival of*Rhopalosiphum padi* (L.) and *Sitobion avenae* (F.)(Homoptera: Aphididae) on different wheat cultivars. Crop Protection, 30(2):216–221, 2011.

2. Gudbjorg I Aradottir and Leonardo Crespo-Herrera. Host plant resistance in wheat to barley yellow dwarf viruses and their aphid vectors: a review. Current Opinion in Insect Science, 45:59–68, 2021.

3. B Bencharki, M El Yamani, and D Zaoui. Assessment of transmission ability of barley yellow dwarf virus-PAV isolates by different populations of *Rhopalosiphum padi* and *Sitobion avenae*. European Journal of Plant Pathology, 106: 455–464, 2000.

4. ME Burrows, MC Caillaud, DM Smith, EC Benson, FE Gildow, and SM Gray. Genetic regulation of polerovirus and luteovirus transmission in the aphid *Schizaphis graminum*. Phytopathology, 96(8):828–837, 2006.

5. ME Burrows, MC Caillaud, DM Smith, and SM Gray. Biometrical genetic analysis of luteovirus transmission in the aphid *Schizaphis graminum*. Heredity, 98(2): 106–113, 2007.

6. S Byrne, M Schughart, V Ballandras, JC Carolan, L Sheppard, and L McNamara. The first survey using high-throughput sequencing of cereal and barley yellow dwarf viruses in Irish spring and winter barley crops. Irish Journal of Agricultural and Food Research, 62:130–145, 2024.

7. Charles-Antoine Dedryver, Anne Le Ralec, and Frédéric Fabre. The conflicting relationships between aphids and men: a review of aphid damage and control strategies. Comptes Rendus Biologies, 333(6-7):539–553, 2010.

8. LC Dempster and SJI Holmes. The incidence of strains of barley yellow dwarf virus in perennial ryegrass crops in south-west and central Scotland. Plant Pathology, 44(4):710–717, 1995.

9. JK Doodson and PJW Saunders. Some effects of barley yellow dwarf virus on spring and winter cereals in field trials. Annals of Applied Biology, 66(3):361–374, 1970.

10. ZQ Du, L Li, L Liu, XF Wang, and G Zhou. Evaluation of aphid transmission abilities and vector transmission phenotypes of barley yellow dwarf viruses in China. Journal of Plant Pathology, pages 251–259, 2007.

11. Séverine Fontaine, Laëtitia Caddoux, and Benoit Barrès. First report of the kdr pyrethroid resistance mutation in a french population of the english grain aphid, sitobion avenae. Crop Protection, 165:106153, 2023.

12. Garth N Foster, Shona Blake, Steve J Tones, Ian Barker, and Richard Harrington. Occurrence of barley yellow dwarf virus in autumn-sown cereal crops in the united kingdom in relation to field characteristics. Pest Management Science: formerly Pesticide Science, 60(2):113–125, 2004.

13. Stephen P Foster, Verity L Paul, Russell Slater, Anne Warren, Ian Denholm, Linda M Field, and Martin S Williamson. A mutation (L1014F) in the voltagegated sodium channel of the grain aphid, *Sitobion avenae*, is associated with resistance to pyrethroid insecticides. Pest management science, 70(8):1249–1253, 2014.

14. Angelica Gonzalez-Gonzalez, Nuri Cabrera, María Eugenia Rubio-Meléndez, Daniela A Sepúlveda, Ricardo Ceballos, Natalí Fernández, Frederic Francis, Christian C Figueroa, and Claudio C Ramirez. Facultative endosymbionts modulate the aphid reproductive performance on wheat cultivars differing in contents of benzoxazinoids. Pest Management Science, 80(4):1949–1956, 2024.

15. Stewart Gray, Michelle Cilia, and Murad Ghanim. Circulative,“nonpropagative” virus transmission: an orchestra of virus-, insect-, and plant-derived instruments. In *Advances in virus research*, volume 89, pages 141–199. Elsevier, 2014.

16. William Gurney and Roger M Nisbet. Ecological dynamics. Oxford University Press, 1998. ISBN 978-0-19-510443-1.

17. Susan E Halbert and KS Pike. Spread of barley yellow dwarf virus and relative importance of local aphid vectors in central Washington. Annals of Applied Biology, 107(3):387–395, 1985.

18. R Harrington, GG Howling, JS Bale, and S Clark. A new approach to the use of meteorological and suction trap data in predicting aphid problems 1. EPPO Bulletin, 21(3):499–505, 1991.

19. MT Howard and AFG Dixon. The effect of plant phenology on the induction of alatae and the development of populations of *Metopolophium dirhodum* (Walker), the rose-grain aphid, on winter wheat. Annals of Applied Biology, 120 (2):203–213, 1992.

20. GUO Jing-Quan, HERVÉ Lapierre, and JEAN-PIERRE Moreau. Vectoring ability of aphid clones of*Rhopalosiphum padi* (L.) and *Sitobion avenae* (Fabr.) and their capacity to retain barley yellow dwarf virus. Annals of applied biology, 131(2):179–188, 1997.

21. DA Kendall, P Brain, and NE Chinn. A simulation model of the epidemiology of barley yellow dwarf virus in winter sown cereals and its application to forecasting. Journal of Applied Ecology, pages 414–426, 1992.

22. Maria Kern, Torsten Meiners, Edgar Schliephake, Antje Habekuss, Frank Ordon, and Torsten Will. Infection of susceptible/tolerant barley genotypes with barley yellow dwarf virus alters the host plant preference of rhopalosiphum padi clones depending upon their ability to transmit bydv. Journal of Pest Science, 95(1): 215–229, 2022.

23. G Lankin-Vega, Sue P Worner, and DAJ Teulon. An ensemble model for predicting *Rhopalosiphum padi abundance*. Entomologia experimentalis et applicata, 129(3): 308–315, 2008.

24. F Leclercq-Le Quillec, Manuel Plantegenest, Gerard Riault, and CA Dedryver. Analyzing and modeling temporal disease progress of barley yellow dwarf virus serotypes in barley fields. Phytopathology, 90(8):860–866, 2000.

25. D Leybourne, M Ramsden, S White, R Wang, H Huang, C Xie, and P Yang. On-line decision support systems, remote sensing and ai applications for wheat pests. Advances in understanding insect pests affecting wheat and other cereals. Burleigh Dodds Science Publishing, UK, pages 411–444, 2023.

26. Daniel J Leybourne. How does diversity within a vector influence yellow dwarf virus transmission efficiency? perspectives from a review. Plant Pathology, 73: 1042–1059, 2024a.

27. Daniel J Leybourne. Cereal aphid fitness: Phenotype data for two cereal aphid species t. 10.17638/datacat.liverpool.ac.uk/2837, 2024b. Accessed: 202410-28.

28. Daniel J Leybourne, Jorunn IB Bos, Tracy A Valentine, and Alison J Karley. The price of protection: a defensive endosymbiont impairs nymph growth in the bird cherry-oat aphid, *Rhopalosiphum padi*. Insect Science, 2018. doi: doi:10.1111/1744-7917.12606.

29. Daniel J Leybourne, Kate E Storer, Abigail Marshall, Nasamu Musa, Samuel Telling, Laurie Abel, Sacha White, Steve Ellis, Po Yang, and Pete M Berry. Thresholds and prediction models to support the sustainable management of herbivorous insects in wheat. a review. Agronomy for Sustainable Development, 44(3):29, 2024a.

30. Daniel J Leybourne, Mark A Whitehead, and Torsten Will. Genetic diversity in vector populations influences the transmission efficiency of an important plant virus. Biology Letters, 20(5):20240095, 2024b.

31. Xiao-Feng Liu, Xiang-Shun Hu, Mike A Keller, Hui-Yan Zhao, Yun-Feng Wu, and Tong-Xian Liu. Tripartite interactions of barley yellow dwarf virus, *Sitobion avenae* and wheat varieties. PLoS One, 9(9):e106639, 2014.

32. Yan Liu, May Oo Khine, Peipei Zhang, Yumei Fu, and Xifeng Wang. Incidence and distribution of insect-transmitted cereal viruses in wheat in china from 2007 to 2019. Plant disease, 104(5):1407–1414, 2020.

33. Leandro Mahieu, Angélica González-González, María Eugenia Rubio-Meléndez, Mario Moya-Hernández, Frederic Francis, and Claudio C Ramírez. An aphid pest superclone benefits from a facultative bacterial endosymbiont in a host-dependent manner, leading to reproductive and proteomic changes. Archives of Insect Biochemistry and Physiology, 117(2):e22154, 2024.

34. Louise McNamara, Kevin Gauthier, Lael Walsh, Gaël Thébaud, Michael Gaffney, and Emmanuel Jacquot. Management of yellow dwarf disease in europe in a post-neonicotinoid agriculture. Pest management science, 76(7):2276–2285, 2020.

35. Derek Morgan. Population dynamics of the bird cherry-oat aphid, *Rhopalosiphum padi* (L.), during the autumn and winter: a modelling approach. Agricultural and Forest Entomology, 2(4):297–304, 2000.

36. K. A. Mottaleb, G. Kruseman, and S. Snapp. Potential impacts of Ukraine-Russia armed conflict on global wheat food security: A quantitative exploration. Global Food Security, 35:100659, 2022.

37. Narelle Nancarrow, Mohammad Aftab, Grant Hollaway, Brendan Rodoni, and Piotr Trębicki. Yield losses caused by barley yellow dwarf virus-PAV infection in wheat and barley: a three-year field study in south-eastern Australia. Microorganisms, 9(3):645, 2021.

38. Keith L Perry, Frederic L Kolb, Bernard Sammons, Clifford Lawson, Gordon Cisar, and Herbert Ohm. Yield effects of barley yellow dwarf virus in soft red winter wheat. Phytopathology, 90(9):1043–1048, 2000.

39. RT Plumb. Barley yellow dwarf virus in aphids caught in suction traps, 1969-73. Annals of Applied Biology, 83(1):53–59, 1976.

40. Katharine F Preedy, Mark AJ Chaplain, Daniel J Leybourne, Glenn Marion, and Alison J Karley. Learning-induced switching costs in a parasitoid can maintain diversity of host aphid phenotypes although biocontrol is destabilized under abiotic stress. Journal of Animal Ecology, 89(5):1216–1229, 2020.

41. FRANÇOISE LECLERCQ-LE QUILLEC, SYLVIE TANGUY, and CA Dedryver. Aerial flow of barley yellow dwarf viruses and of their vectors in western France. Annals of applied Biology, 126(1):75–90, 1995.

42. R Rabbinge, EM Drees, M Van der Graaf, FCM Verberne, and A Wesselo. Damage effects of cereal aphids in wheat. Netherlands Journal of Plant Pathology, 87: 217–232, 1981.

43. Guillermo E Ramírez-Cáceres, Mario G Moya-Hernández, Manuel Quilodrán, Roberto F Nespolo, Ricardo Ceballos, Cristian A Villagra, and Claudio C Ramírez. Harbouring the secondary endosymbiont *Regiella insecticola* increases predation risk and reproduction in the cereal aphid sitobion avenae. Journal of Pest Science, 92:1039–1047, 2019.

44. WF Rochow and VF Eastop. Variation within *Rhopalosiphum padi* and transmission of barley yellow dwarf virus by clones of four aphid species. Virology, 30 (2):286–296, 1966.

45. CD Rogers, RML Guimarães, KA Evans, and SA Rogers. Spatial and temporal analysis of wheat bulb fly (*Delia coarctata*, Fallén) oviposition: consequences for pest population monitoring. Journal of pest science, 88:75–86, 2015.

46. Patricia Sanches, Consuelo M De Moraes, and Mark C Mescher. Endosymbionts modulate virus effects on aphid-plant interactions. The ISME Journal, 17(12): 2441–2451, 2023.

47. DJ Thackray, AJ Diggle, and RAC Jones. BYDV PREDICTOR: a simulation model to predict aphid arrival, epidemics of barley yellow dwarf virus and yield losses in wheat crops in a mediterranean-type environment. Plant Pathology, 58 (1):186–202, 2009.

48. Lael E Walsh, Olaf Schmidt, Martin S Williamson, and Michael T Gaffney. Infield prevalence of resistant grain aphid *Sitobion avenae* (Fabricius). In *Biology and Environment: Proceedings of the Royal Irish Academy*, volume 120, pages 29–38. JSTOR, 2020.

49. AD Watt. The effect of cereal growth stages on the reproductive activity of *Sitobion avenae* and *Metopolophium dirhodum*. Annals of Applied Biology, 91(2):147–157, 1979.

50. Sacha White, Sam Telling, Hannah Griffiths, Dave Skirvin, M Williamson, S A Ellis, Tim Schaare, R Granger, and O Potter. Management of aphid and BYDV risk in winter cereals. AHDB Project Report, 646:1–157, 2023.

51. Chenchen Zhao and Meixue Zhou. Effect of barley yellow dwarf virus (BYDV) on barley: A precise assessment of reductions in yield components under variable disease severities. Plant Disease, 2024. doi: 10.1094/PDIS-04-24-0883-SC.

52. Sharon E Zytynska, Karim Tighiouart, and Enric Frago. Benefits and costs of hosting facultative symbionts in plant-sucking insects: A meta-analysis. Molecular Ecology, 30(11):2483–2494, 2021.

